# Characterization of novel cytoplasmic roles for the N-terminal methyltransferase NRMT1

**DOI:** 10.64898/2026.01.17.700099

**Authors:** John G. Tooley, Gao Zhou, Sarah Obeidat, Arielle Arbel, Cameron Jones, Frank Tedeschi, Christine E. Schaner Tooley

## Abstract

N-terminal methylation of proteins by the trimethylase NRMT1 plays important roles in oncogenesis, development, and aging. As N-terminal methylation has frequently been shown to regulate protein-DNA interactions, and many NRMT1 substrates are transcription factors or regulators of chromatin structure, previous research has focused on how transcriptional regulation by NRMT1 affects cell growth and differentiation. However, we have recently identified a new, cytoplasmic role for NRMT1, inhibiting the eukaryotic elongation factor 1 alpha (eEF1A) methyltransferase METTL13, which indicates NRMT1 could also be acting as a translational regulator. Here we further explore NRMT1 cytoplasmic functions and show that, unlike previously thought, NRMT1 can methylate substrates in the cytoplasm. We also show that while many of these substrates remain bound to NRMT1, it can also interact with a number of non-target ribosomal proteins and proteins associated with the endoplasmic reticulum (ER). To confirm NRMT1 interaction with the ribosome, we performed polysome profiling, which showed a portion of NRMT1 co-migrates with the 40S and 60S subunits but not with actively translating polysomes, indicating NRMT1 may play an early role in translation. To see if NRMT1 was affecting target mRNA selection of ribosomes, we also performed ribosome-sequencing experiments in proliferating and differentiating C2C12 mouse myoblasts. These results show a striking upregulation of translation of soluble proteins with NRMT1 loss and corresponding decrease in translation of transmembrane and signal sequence-containing proteins. We now propose a model where NRMT1 regulates the translation of transmembrane and secreted proteins by facilitating interactions between the ribosome and the ER.

## Introduction

The first eukaryotic N-terminal methyltransferase, NRMT1 (also known as METTL11A and NTMT1), was discovered over 15 years ago^1,2^. It was originally thought to exclusively methylate substrates that contain an N-terminal X-Pro-Lys consensus sequence after cleavage of the initiating methionine^3^. However, after its discovery it was also shown to methylate a non-canonical consensus that could include Ala/Gly/Ser in the second position and Arg in the third position^4^. This expanded consensus sequence indicates NRMT1 has over 300 potential substrates, many of which are transcription factors or regulators of chromatin structure^1,4^.

Given the known functions of many of its substrates and the fact that N-terminal trimethylation results in a pH-insensitive positive charge on the N-terminus, it was hypothesized that N-terminal methylation may regulate protein interaction with the negative DNA backbone^5^. This was shown to be true for the NRMT1 substrates RCC1, DDB2, CENP-A, CENP-B, MYL9, and ZHX2^6–11^. Accordingly, NRMT1 loss results in defects in mitosis, DNA repair, and transcriptional regulation^1,6,7,10,11^, and leads to oncogenic growth and defects in stem cell differentiation^12–14^.

As many of the first NRMT1 targets to be characterized performed functions in the nucleus, and NRMT1 was originally shown to primarily localize to and be active in the nucleus^1,4^, it was initially thought to be a nuclear methyltransferase. However, in budding yeast NRMT1 was originally identified in a screen for regulators of protein translation^15^, and our recent work demonstrated that NRMT1 can also localize to the cytoplasm and regulate the activity of the eukaryotic elongation factor 1 alpha (eEF1A) methyltransferase METTL13^16^. Additionally, many canonical NRMT1 targets are predominantly localized to the cytoplasm, including cytoskeletal proteins (MYL1, MYL2, MYL6B, MYL11), ER-associated proteins (VRK2, SSR3), and ribosomal proteins (RPS25, RPL12, RPL23A)^1,2,4^.

In this study, we aimed to better understand the function of NRMT1 in the cytoplasm, specifically focusing on its role in translational regulation. We found that NRMT1 localizes to both the soluble cytosol and membrane-bound organelle fractions of the cytoplasm and that cytoplasmic NRMT1 has substrate-specific methylation activity. We also found that, in addition to its substrates, NRMT1 interacts with numerous non-target ribosomal proteins (RPs), ribosomal binding proteins (RBPs), and proteins associated with the endoplasmic reticulum (ER). Polysome profiling indicated NRMT1 does not bind to actively translating ribosomes but does bind individual RPs and co-migrates most strongly with the 40S and 60S ribosomal subunits. Finally, ribosome-sequencing showed that loss of NRMT1 in proliferating and differentiating C2C12 cells results in increased translation of soluble proteins and decreased translation of transmembrane and signal sequence-containing proteins, including those involved in chemotaxis. These data extend the impact of N-terminal methylation outside of the previously ascribed nuclear functions and support a model where NRMT1 helps direct stem cell differentiation through facilitating interactions between the ribosome and the ER.

## Results

### Cytoplasmic localization and activity of NRMT1

We have previously shown by immunofluorescence that transfected GFP-NRMT1 appears primarily nuclear^1^ and inactive toward a cytosolically expressed RCC1 peptide^4^. However, more recent cellular fractionation of HEK293T cells revealed that NRMT1 also localizes to the cytoplasm^16^. To determine if this localization pattern extends to other cell types, we performed nuclear/cytoplasmic fractionations of the HeLa cervical carcinoma and C2C12 mouse myoblast cell lines. As is seen for HEK293T cells, NRMT1 also localizes to both the nucleus and the cytoplasm in HeLa and C2C12 cells, with more NRMT1 in the cytoplasm of both (**Fig. 1A**). We also found that NRMT1 localization patterns are not significantly disrupted by treatment with the nuclear export inhibitor Leptomycin B, suggesting that the NRMT1 enzyme (predicted 26 kDa) may be able to passively diffuse the nuclear pore (**Fig. 1B**). Additional sub fractionation of cytosolic extracts revealed that NRMT1 localizes to both the soluble cytosol, as well as membrane-bound organelles, including the ER (**Fig. 1C**).

**Fig. 1 –.**
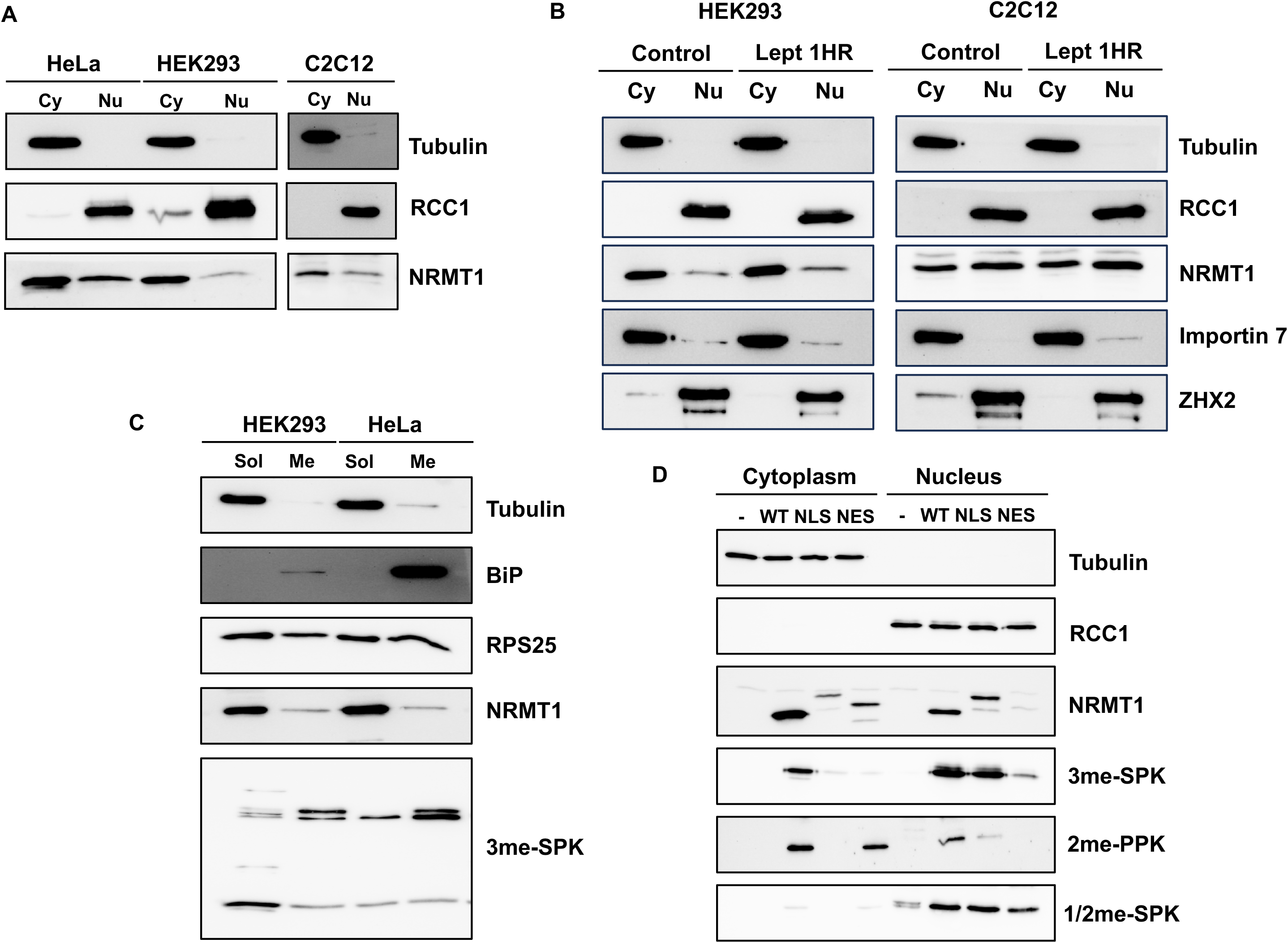
NRMT1 localizes to the cytoplasm in numerous cell lines and has substrate-specific cytoplasmic activity. Western blots showing **A**) NRMT1 localizes to the nuclear (Nu) and cytoplasmic (Cy) fractions in HeLa, HEK293, and C2C12 cells. Tubulin (cytoplasmic) and RCC1 (nuclear) are used for fractionation controls. **B**) 1 hour of Leptomycin B (Lept) treatment does not alter NRMT1 localization patterns in HEK293 or C2C12 cells, indicating it could be passively diffusing through the nuclear pore. Importin 7 and ZHX2 are controls for Leptomycin B function. **C**) NRMT1 and its methylated targets (3me-SPK) are present in both the soluble and membrane-bound cytoplasmic fractions. BiP is a control for the membrane-bound fraction. Tubulin is a control for the soluble fraction, while RPS25 is present in both. **D**) Wild type (WT) NRMT1 can increase all three types of methylation (3me-SPK, 2me-SPK, and 1/2me-SPK) in the cytoplasm and nuclear fractions of NRMT1 KO HEK293 cells. NRMT1 fused to a nuclear localization signal (NLS) predominantly restores methylation to SPK targets in the nucleus, while NRMT1 fused to a nuclear export signal (NES) can restore 2me-PPK methylation in the cytoplasm.

As we have previously seen that fusing a nuclear export signal (NES) to an RCC1-GFP fusion protein (SPK consensus) inhibits its trimethylation^4^, we next asked if we could detect N-terminal methylation of any cytoplasmic targets. To test this, we fused either a nuclear localization signal (NLS) or NES onto NRMT1 to localize it to a specific compartment. In addition, a 2xGFP tag was added to increase the size of the protein and prevent passive diffusion through the nuclear pore. Constructs were expressed in HEK293 NRMT1 knockout (KO) cells, which lack both NRMT1 expression and fully methylated target proteins (**Fig. 1D**). We can detect low levels of mono/di-methylated products in the nucleus of this cell line, but we attribute this to the presence of NRMT2 ^17,18^. In cells expressing WT-NRMT1-GFP, we see some restoration of all types of methylated targets in both cellular compartments (**Fig.1D**). In cells expressing NLS-NRMT1-GFP, even though there is some residual NRMT1 expression in the cytosol, restoration of methylated targets was only seen in the nucleus, and this was target specific. Methylation of the targets recognized by the 3me-SPK and 1/2me-SPK antibodies was restored similar to what was seen with WT NRMT1. However, the target recognized by the 2me-PPK antibody was only slightly restored (**Fig. 1D**). Conversely, in cells expressing NES- NRMT1-GFP, the target recognized by the 2me-PPK antibody is the only one whose methylation is restored in the cytoplasm (**Fig. 1D**). These results suggest that NRMT1 is active in the cytoplasm, but this activity is dependent on the consensus sequence of the target.

### Cytoplasmic interactors of NRMT1

We have recently identified a cytoplasmic interaction between NRMT1 and its family member METTL13 and went on to show that this interaction negatively regulates NRMT1 activity^16^. To identify additional novel cytoplasmic interactors of NRMT1, we performed cytoplasmic/nuclear fractionation of HEK293T cells transfected with NRMT1-FLAG. NRMT1-FLAG was immunoprecipitated from the cytoplasmic fraction, and liquid chromatography-mass spectrometry was performed to identify interacting proteins. Our first observation was that NRMT1 interacted with multiple targets, both canonical and non-canonical, many of which were identified as having a 100% abundance ratio as compared to untransfected control cells (**Fig. 2A**). This was unsurprising, as NRMT1 has previously been shown to tightly bind its methylated products^4^.

**Fig. 2 –.**
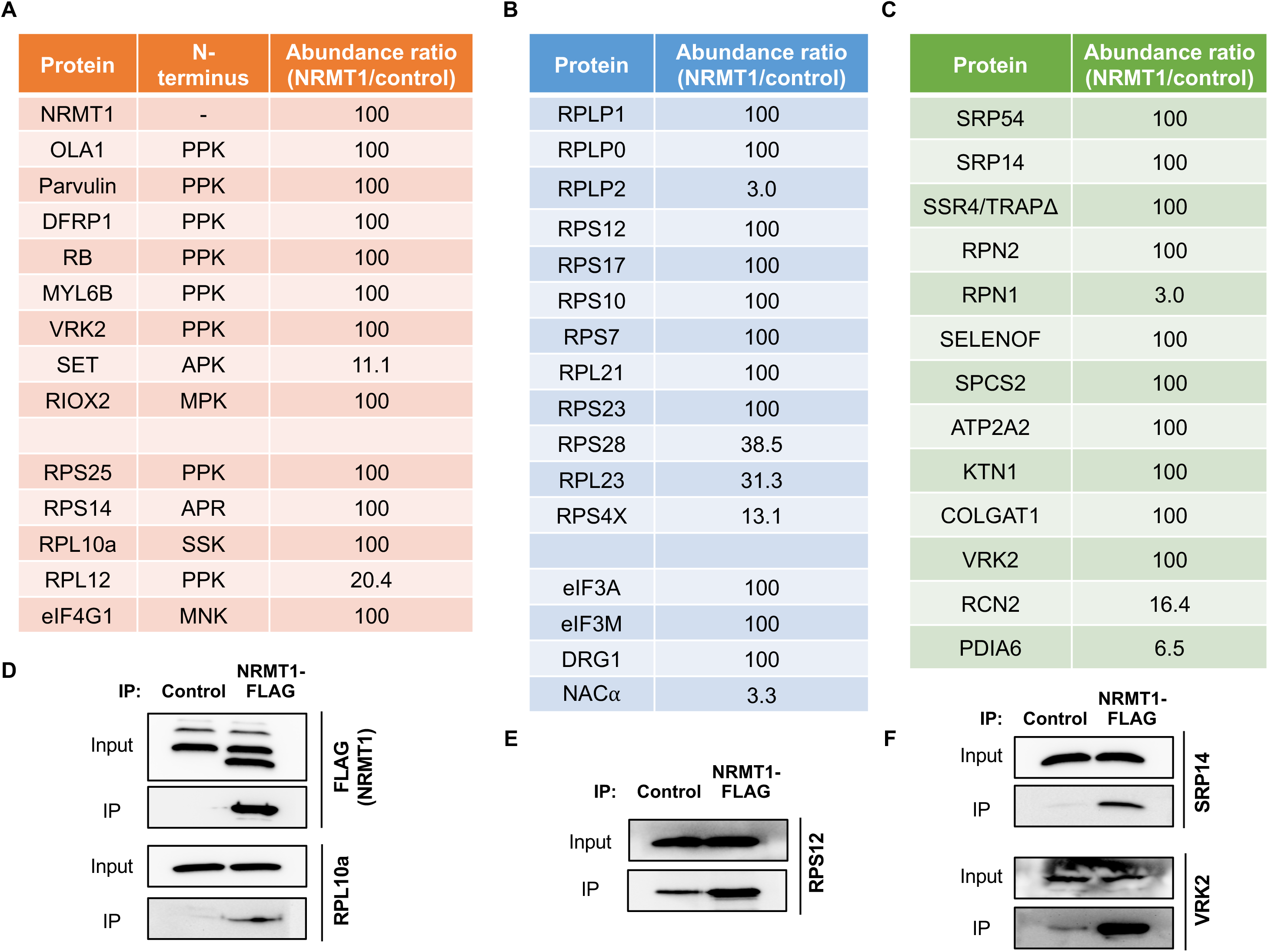
Cytoplasmic interactors of NRMT1 identified by MS analysis. In the cytoplasm, NRMT1 interacts with **A**) ribosomal and non-ribosomal targets, many of which contain the PPK consensus, **B**) non-target ribosomal proteins and other proteins involved in translation (lower section), and **C**) proteins associated with the ER. **D-F**) Western blots verifying NRMT-FLAG co-IPs with selected targets, RPL10a, RPS12, VRK2, and SRP14.

Our next observation was that there was a significant enrichment of non-target ribosomal proteins (**Fig. 2B**). Components of both the large and small subunits were represented, and often the individual RPs identified localized to the interaction interface between the two subunits. Additionally, all members of the transiently associated P-stalk complex were enriched in NRMT1 pulldowns, including the NRMT1 substrate RPL12, which serves as the base for the P-stalk on the ribosomal large subunit. Interestingly, we also detected several proteins known to localize to the ER (**Fig. 2C**). These include the NRMT1 target VRK2, the signal recognition particle complex members SRP54 and SRP14, the translocon-associated protein (TRAP) complex member SSR4, the Oligosaccharyltransferase (OST) complex members RPN1 and RPN2, and the signal peptidase complex (SPC) member SPCS2 (**Fig. 2C**). In addition, there were a variety of cytoplasmic NRMT1 interacting proteins that have a more general association with protein translation (**Fig. 2A,B**). These include the NRMT1 targets OLA1, Parvulin, RIOX2, and DFRP1/ZC3H15 ^19–22^, as well as the DFRP1 binding partner DRG1. We also identified interactions with eukaryotic initiation factors (eIFs), NACα and the RNA helicase DDX3X – all of which have been shown to bind ribosomes or selectively impact translation^23–25^. Representatives from each group were confirmed through co-immunoprecipitations with NRMT1-FLAG (**Fig. 2D-F**).

### Role of NRMT1 in translation

One of the first phenotypes ascribed to NRMT1 came from a screen of the *S.* cerevisiae non-essential gene deletion array (yGDA) for novel proteins that caused translational defects^15^. Previously an uncharacterized gene, its loss was found to alter the ribosome profile and cause defects in translation efficiency and fidelity, and it was given the name translation associated element 1 (TAE1)^15^. As such, we set out to further examine the role of NRMT1 in mammalian protein translation. We first asked if loss of NRMT1 caused cellular stress that led to an overall reduction of translation levels, as evidenced by induction of the Integrated Stress Response (ISR) and phosphorylation of eIF2α. We found that loss of NRMT1 did not significantly alter ISR induction, either in basal culture conditions or in response to cellular stressors (**Fig. 3A**). While RPs are predominantly cytoplasmic, they are temporarily transported into the nucleolus to be assembled into pre-ribosomal subunits before returning to the cytosol ^26^. To determine if NRMT1 is playing a role in ribosome biogenesis, we performed sucrose gradient analysis of nuclear fractions and found that NRMT1 failed to co-migrate with either small or large pre-ribosomal subunits (**Fig. 3B**). Additionally, GFP-NRMT1 transfected in HEK293 cells failed to localize to the nucleolus (data not shown), indicating that NRMT1 does not play a role in ribosome biogenesis.

**Fig. 3 –.**
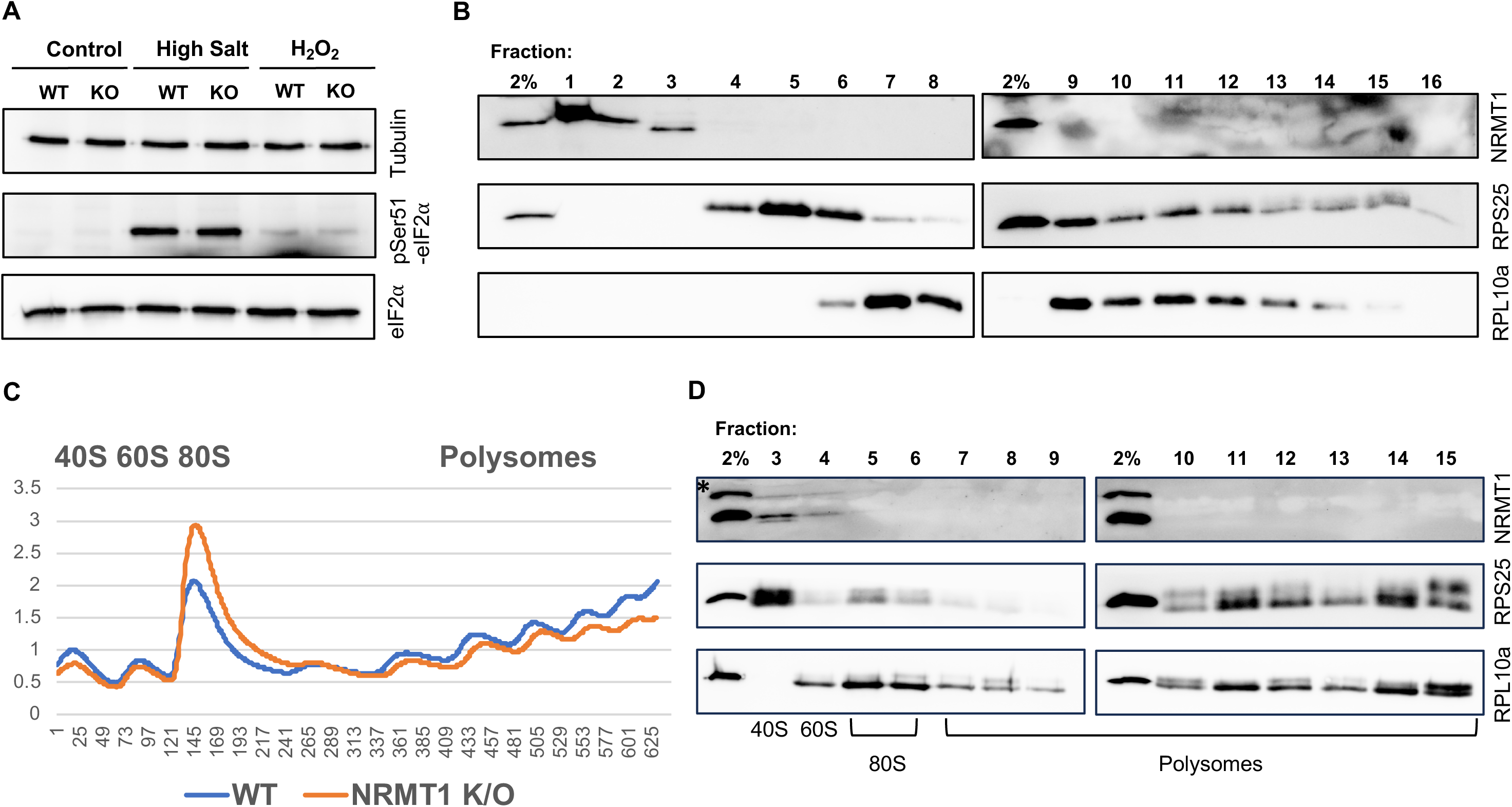
NRMT1 is not involved in the integrated stress response or ribosome biogenesis but does co-migrate with the 40S and 60S subunits. **A**) Western blot showing knockout (KO) of NRMT1 does not increase levels of eIF2α phosphorylated at Ser 51 (pSer51-eIF2α) even under conditions of high salt or hydrogen peroxide (H_2_O_2_). **B**) Sucrose gradient analysis of nuclear fractions showed it also fails to co-migrate with either small or large pre-ribosomal subunits, as indicated by exclusion from fractions containing RPS25 and RPL10A, respectively. **C, D**) Polysome profile UV trace and corresponding Western blot analysis in HEK293 cells showing NRMT co-migrates most strongly with the 40S and 60S ribosomal subunits, and its loss results in increased monosome formation and corresponding decreased polysome formation.

Newly generated proteins can be N-terminally modified as soon as a nascent peptide emerges from the exit tunnel of the ribosome, or post-translationally following protein synthesis ^27^. Since NRMT1 localizes to the cytoplasm and binds many RPs and RBPs, we wondered if NRMT1 could be modifying proteins co-translationally, similar to what is seen for the N-terminal acetyltransferase NatA and the N-terminal myristoyltransferase (NMT) complexes ^28–31^. We used polysome profiling in HEK293 cells to see if NRMT1 co-migrated with any ribosomal fractions and found that instead of co-sedimenting with actively translating polysomes, NRMT1 most strongly co-migrated with the 40S and 60S subunits (**Fig. 3C**), as indicating by probing the sucrose gradient fractions for NRMT1, the 40S subunit protein RPS25, and the 60S subunit protein RPL10a (**Fig.3D**). Additionally, there was an increase in 80S monosomes in the KO cells and a corresponding decrease in polysomes (**Fig. 3C**). From these data, we conclude that NRMT1 interactions with RPs and RBPs largely occur with individual proteins and ribosomal subunits and not on polysomes.

### Role of NRMT1 in ribosomal transcript selection

We had previously seen that loss of NRMT1 in C2C12 cells impairs their differentiation into myotubes^14^. To understand if NRMT1 was altering the muscle-specific transcriptional profile, we then performed RNA-sequencing of WT and NRMT1 KO C2C12 cells under basal growth conditions (day 0) and after one day of differentiation (day 1). Surprisingly, at day 1, we found that the KO cells were able to upregulate most of the muscle-specific transcriptional program, similar to WT cells^32^. However, transcription of secreted proteins, including those involved in chemotaxis was mis-regulated. To now determine if there was also an alteration in the translational regulation of these cells, we performed ribosome profiling (Ribo-Seq) under the same conditions.

Ribo-Seq provides a snapshot of the identities and levels of mRNAs being translated at a selected time point. Each transcript detected is denoted as a ribosome protected fragment (RPF). At day 0 (proliferating, undifferentiated myoblasts), RPFs were generated for 10,348 individual genes common to both WT and KO cells. We first used Cytoscape to perform Gene Ontology (GO) analysis of the top 50 most-downregulated RPFs in the NRMT1 KO cells at this time point. This functional enrichment analysis uncovered a significant enrichment in both GO cellular compartment (ER) and molecular function (Growth Factor Binding). A manual examination of these proteins in the UniProt database confirmed that 70% of the proteins identified contained a signal sequence (SS) and/or transmembrane domain (TMD) (**Fig. 4A**), a percentage that is significantly higher than the ~30% of the overall proteome that traverses the ER^33^. Additionally, eight of the proteins identified as having a transmembrane domain localized to the ER membrane, and another 4 of the 50 proteins identified were ribosomal proteins (**Fig. 4A**). We verified two of the hits that contain both a SS and TMD by Western blot, growth hormone receptor (GHR) and leukemia inhibitory factor receptor (LIFR). The higher molecular weight bands of each, reflecting the mature, glycosylated version, were reduced in NRMT1 KO cells (**Fig. 4B**).

**Fig. 4 –.**
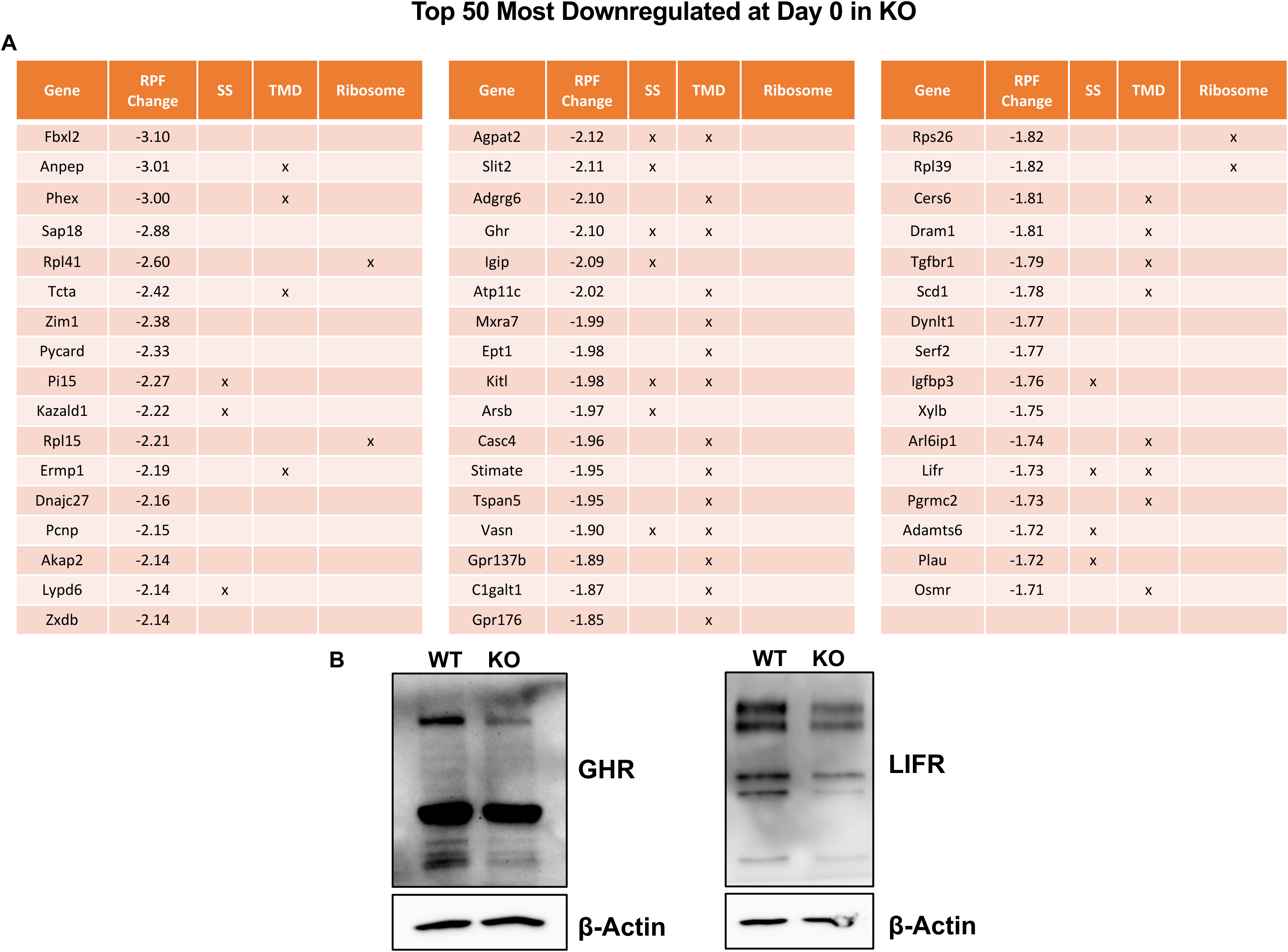
Loss of NRMT1 in proliferating C2C12 cells decreases translation of signal sequence-containing, transmembrane, and ribosomal proteins. **A**) Top 50 mRNAs with downregulated translation at day 0, as quantified by change in ribosome protected fragments (RPFs). 70% of the proteins identified contained a signal sequence (SS) and/or transmembrane domain (TMD). Four were also ribosomal proteins. **B**) Western blot confirmation of decreased levels of growth hormone receptor (GHR) and leukemia inhibitory factor receptor (LIFR) in NRMT1 knockout (KO) C2C12 myoblasts as compared to wild type (WT).

We observed a similar pattern when examining the top 50 mRNAs whose RPF levels were downregulated at day 1 in NRMT1 KO cells. Here, 58% of the proteins contained a SS and/or TMD, and three of the mRNAs identified were captured at both days 0 and 1 (**Fig. 5**). These included the mRNAs that encode for DYNLT1, an accessory to the dynein complex that can localize to Golgi^34^; ARL6IP1, an ER transmembrane protein which plays a role in the formation and stabilization of ER tubules^35^; and the above-mentioned GHR, a transmembrane protein that regulates postnatal body growth and in humans can also be cleaved to generate a soluble peptide^36^. Taken together, these data suggest that translation of secreted and ER-associated proteins is impaired by NRMT1 loss.

**Fig. 5 –.**
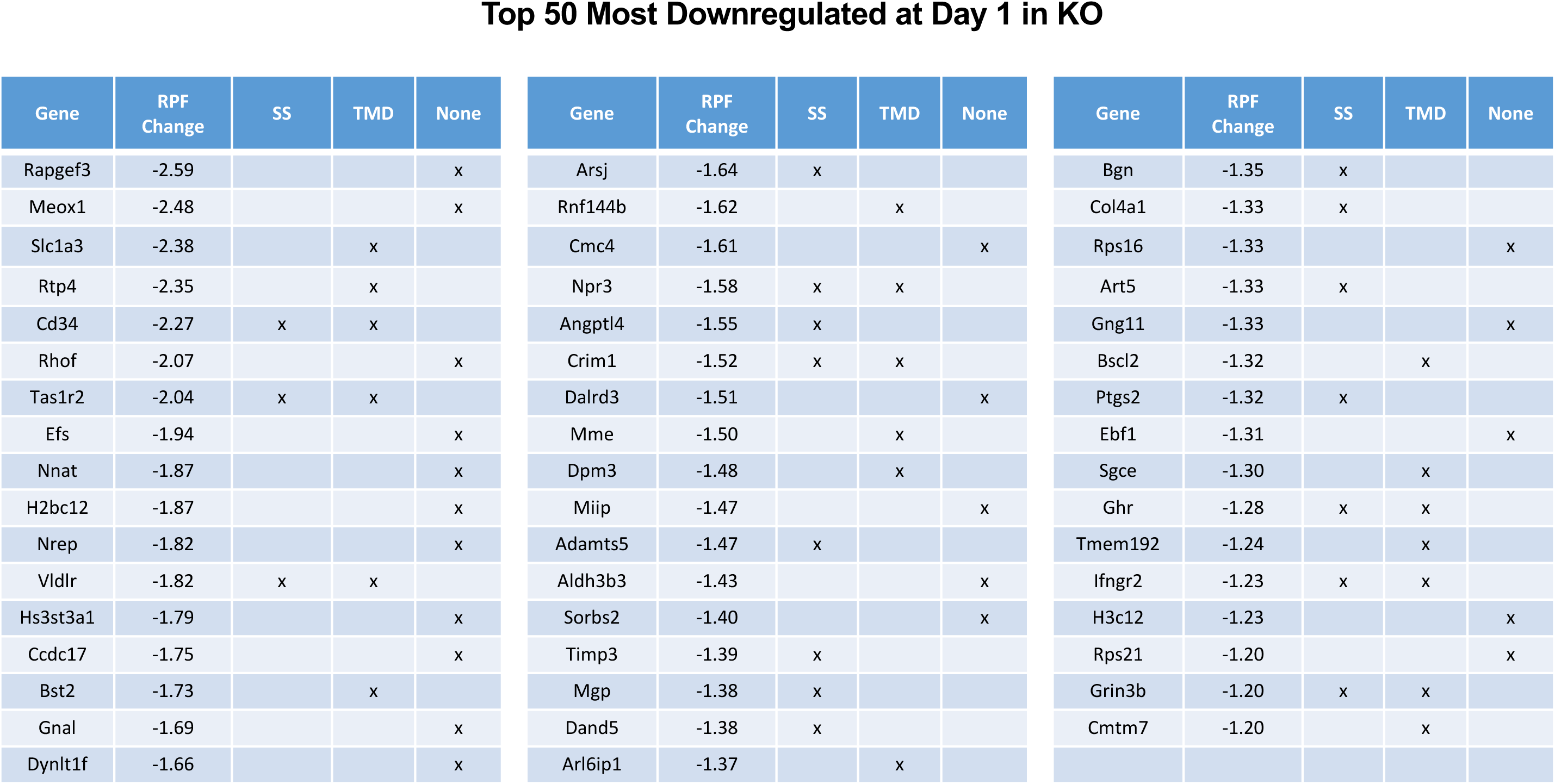
Loss of NRMT1 in differentiating C2C12 cells decreases translation of signal sequence-containing and transmembrane proteins. Top 50 mRNAs with downregulated translation at day 1, as quantified by change in RPFs. 58% of the proteins contained a SS and/or TMD. Three mRNAs were in the top 50 for both days 0 and 1, *Dynlt1*, *Arl6ip1*, and *Ghr*.

In contrast, mRNAs whose RPF levels were upregulated in NRMT1 KO cells showed none of the enrichment for secretory or transmembrane properties. Rather, evaluation of the top 50 mRNAs with upregulated RPFs showed a disproportionally lower percentage of ER-traversing proteins, both at days 0 and 1 (**Figs. 6 and 7**). Only 8% (day 0) and 14% (day 1) of proteins evaluated contained a SS and/or TMD, well below the estimated 30% of the overall proteome. Again, three mRNAs showed increased translation at both time points. These included *Bbs7*, which encodes a member of the Bbsome complex involved in cilia formation; *Gin1*, which encodes a DNA binding protein involved in DNA integration; and *Pinx1*, which encodes a protein involved in telomere maintenance. BBS7 is predominantly cytosolic, while GIN1 and PINX1 are normally found in the nucleus or nucleoplasm^37–39^.

**Fig. 6 –.**
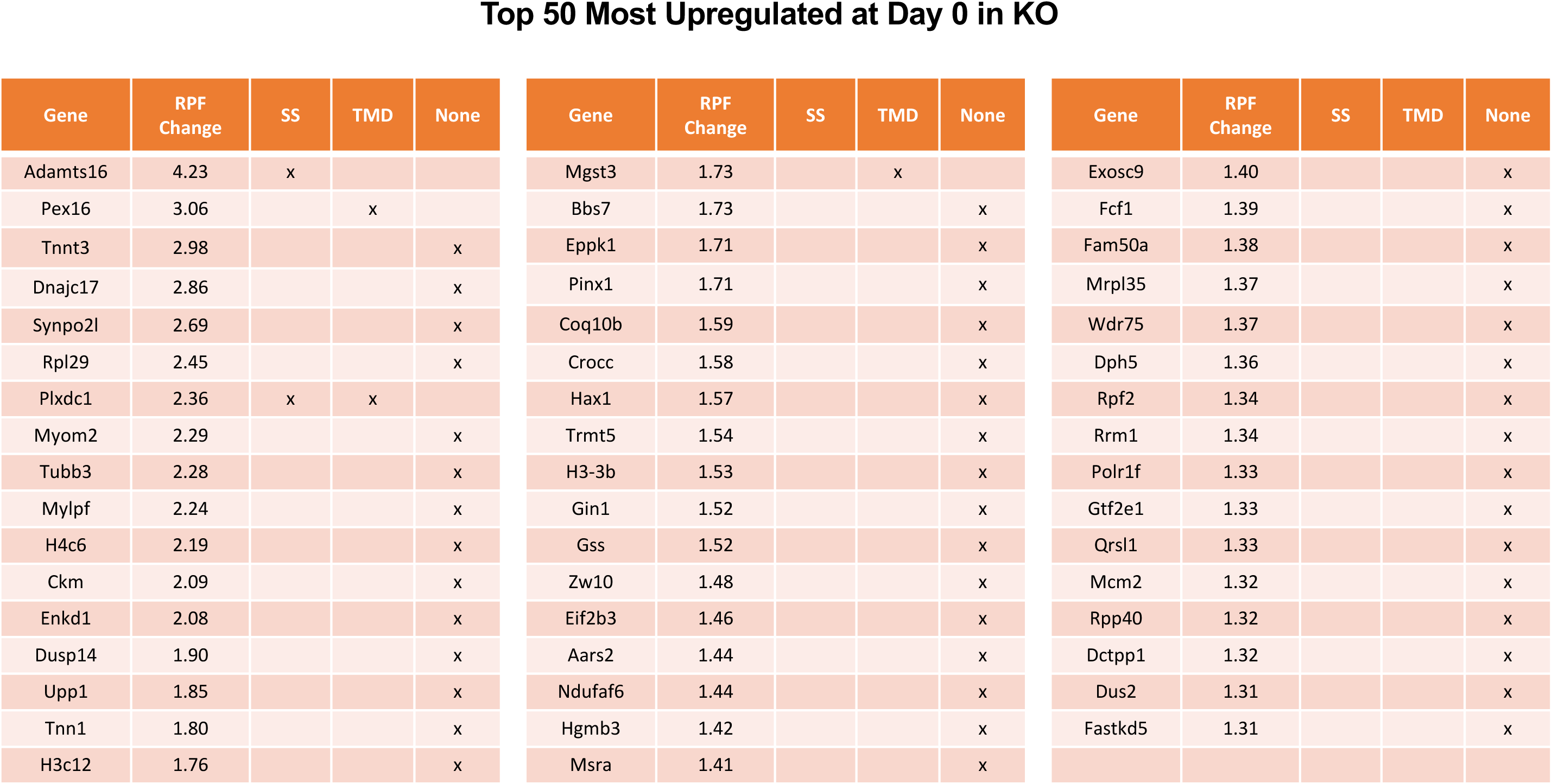
Loss of NRMT1 in proliferating C2C12 cells increases translation of soluble proteins. Top 50 mRNAs with upregulated translation at day 0, as quantified by change in RPFs. Only 8% of proteins evaluated contained a SS and/or TMD, well below the estimated 30% of the overall proteome.

**Fig. 7 –.**
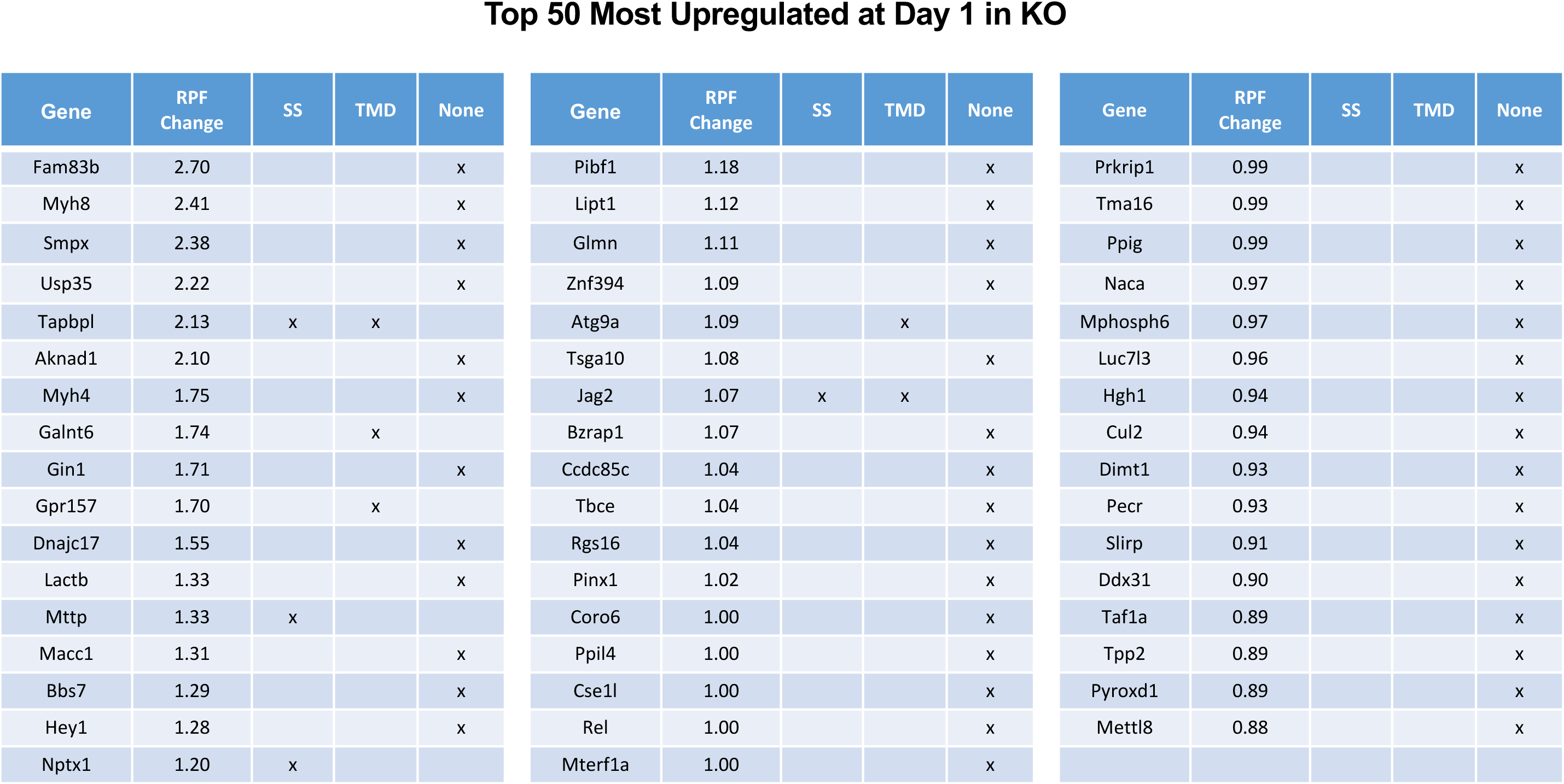
Loss of NRMT1 in differentiating C2C12 cells increases translation of soluble proteins. Top 50 mRNAs with upregulated translation at day1, as quantified by change in RPFs. Only 14% of proteins evaluated contained a SS and/or TMD. Three mRNAs were in the top 50 for both day 0 and day1, *Bbs7*, *Gin1*, and *Pinx1*.

## Discussion

These data confirm our previous studies showing a portion of NRMT1 localizes to the cytoplasm^16^. This was surprising, as we had also previously shown that an RCC1-GFP fusion protein with an SPK consensus sequence and NES could not be trimethylated in the cytoplasm^4^. Here we explain this, by showing that cytoplasmic Nα-methylation is consensus sequence-specific (**Fig. 1D**). WT NRMT1 could methylate both SPK and PPK substrates in the nucleus and cytoplasm, while the NLS-containing NRMT1 could only match WT methylation levels of the SPK substrates in the nucleus. Conversely, the NES-containing NRMT1 could only match WT methylation levels of the PPK substrate in the cytoplasm. This PPK-specific cytoplasmic activity of NRMT1 makes sense considering many of the ribosomal targets and other cytoplasmic targets contain this consensus sequence (**Fig. 2A**)^4^.

Though the N-terminal sequences of many ribosomal proteins have evolved, Nα-methylation of RPs is an ancient and conserved process that was first described decades ago^40^. NRMT1 has been shown to bind its methylated targets tightly^4^, and our NRMT1 pulldowns were highly enriched for both target and non-target RPs, leading us to wonder if NRMT1 was capable of binding polysomes, via its methylated products, and co-translationally modifying nascent protein chains. This is a well-established function of other N-terminal protein modifications, including acetylation and myristylation. The ribosome is a hub of transient, sometimes competitive interactions by protein modifying enzymes, yet only a fraction of total ribosomes are occupied at any given time by many of these proteins^41,42,43,44^. Therefore, it seemed possible that NRMT1 itself was capable of binding a subset of actively translating ribosomes and acting co-translationally, which would generate a pool of specialized ribosomes endowed with unique regulatory capabilities^45^.

However, our polysome profiling shows that NRMT1 does not localize to polysomes but rather co-migrates with the 40S and 60S subunits, indicating co-translational Nα-methylation is not likely occurring. Alternatively, our NRMT1 pulldown experiments uncovered interactions with the eIF3A and eIF3M proteins, which transiently associate with the 40S subunit and facilitate pre-initiation complex formation, mRNA scanning, and the prevention of premature joining of the 40S and 60S subunits^46^. NRMT1 also interacts strongly with its target OLA1, which can bind the 60S subunit and split the subunits of stalled ribosomes^47^, as well as target protein DFRP1, which can bind RPS12 on the 40S subunit and brings its binding partner DRG1 to stalled ribosomes, where its promotes a favorable conformation to allow translation to proceed^22^. Our pulldown experiments also captured all components of the P stalk complex, which specializes in translating secreted cytokines^48^ and TMD-containing proteins^49^ and may help reduce the stalls inherently induced by TMDs during translation. Notably, all these NRMT1 interactors bind the ribosome transiently and often only under certain conditions, suggesting that any interaction of NRMT1 with the translational machinery might also be transient and situation-based.

Our pulldown experiments also detected interactions between NRMT1 and the SRP complex members SRP54 and SRP14. Nascent peptides destined for the ER first encounter the SRP at the ribosome peptide exit tunnel, and we did capture slight interactions with both NACα and its ribosome anchor RPL23 (**Fig. 2B**). NAC normally antagonizes SRP binding, unless it encounters a peptide with an ER targeting sequence, and it is tempting to speculate that we are capturing a subset of ribosomes translating proteins destined for the ER. Notably, we did not capture any interactions with the NatA complex or the NMTs. Upon binding of a SS by SRP, translation is paused by the Alu domain of SRP (which includes SRP14), which allows the ribosome to translocate to the ER and properly insert the nascent peptide into the membrane before translation resumes. This mechanism prevents premature translation and improper folding of the protein in the soluble cytosol. Taken together, these data suggest a model where NRMT1 interacts transiently with the early ribosome, and when a SS peptide becomes exposed, helps facilitate the delivery of that ribosome to the ER. Whether this is a catalytic or non-catalytic function of NRMT1 remains to be determined.

NRMT1 plays important roles in oncogenesis and stem cell development^12–14^, and until now, mechanistic studies have primarily focused on how its role in transcriptional regulation resulted in loss of function phenotypes^10,11^. Here, we show that NRMT1 not only also localizes to the cytoplasm, but also has substrate-specific cytoplasmic activity, which could promote its role as a facilitator of SS-containing ribosome transport to the ER. These data suggest a new role for NRMT1 in translational regulation of secreted proteins that could contribute to the oncogenic and stem cell differentiation phenotypes seen with NRMT1 loss. For example, the decrease in GHR translation in NRMT1 KO myoblasts could contribute to the observed differentiation defects by inhibiting migration and fusion, as previously seen in poultry muscle cells^50^. Similarly, decreased translation of DYNLT1 could also contribute to impaired myoblast differentiation, as the dynein complex is needed for nuclear movement during myotube formation^51^. Translational control of stem cell differentiation is an emerging field of study, and a better mechanistic understanding will increase the therapeutic options for treatment of numerous degenerative disorders.

## Materials and methods

### Cell culture and treatments

Human embryonic kidney (HEK-293T), human cervical cancer (HeLa), and C2C12 mouse myoblasts were cultured in Dulbecco’s Modified Eagle Medium (DMEM; ThermoFisher, Grand Island, NY) with 10% fetal bovine serum (FBS; Atlanta Biologicals, Atlanta, GA) and 1% penicillin-streptomycin (P/S; ThermoFisher). All cells were grown at 37°C and 5% CO_2_ on tissue culture treated plastic (Corning, Corning, NY). C2C12 cells were a generous gift from Dr. Ian Macara, Vanderbilt University. For experiments involving C2C12 muscle cell differentiation, cells were washed with PBS and then incubated in differentiation media (DMEM, 2% horse serum, 50 nM insulin (Sigma, St. Louis, MO)) for 24 hours. For cell stress experiments, cells were treated for 1 hour with either full DMEM media supplemented with 500 mM NaCl (High Salt), or 1 mM H_2_O_2_. For experiments to impair nuclear export, cells were treated with 10 ng/mL (18.5 nM, in EtOH) Leptomycin B (Cayman Chemical, Ann Arbor, MI) for 1 hour prior to harvest.

### Cellular fractionations

To generate nuclear/cytoplasmic fractions, cells were washed with PBS, collected by trypsinization, and resuspended into cytoplasmic lysis buffer (10 mM HEPES, 10 mM KCl, 1.5 mM MgCl_2_, 0.5 mM DTT, 0.3% NP-40, pH 7.9) plus protease inhibitors. Lysates were incubated on ice for 25 min with occasional vortexing, then spun down at 2,500 rpm for 4 min at 4°C. Supernatant was retrieved as cytoplasmic fraction. Nuclear pellet was thoroughly washed in cytoplasmic lysis buffer by three cycles of resuspension and centrifugation. Following final wash step, pellet was resuspended in nuclear extraction buffer (20 mM HEPES, 450 mM NaCl, 1 mM EDTA, 0.5 mM DTT, 26% glycerol, pH 7.9) plus protease inhibitors. Sample was incubated on ice for 30 min. with vortexing every 10 min. Lysate was then spun down at 12,000 rpm for 10 min at 4°C, and supernatant was collected as the nuclear fraction.

To sub-fractionate cytoplasmic extracts into soluble and membrane-bound fractions, cells grown on 10-cm plates were initially washed with ice-cold PBS. After all liquid was thoroughly aspirated, the plate was gently coated with 0.5 mL of permeabilization buffer (110 mM KOAc, 25 mM K-HEPES, 2.5 mM Mg(OAc)_2_, 1mM EGTA, 0.015% Digitonin, 1 mM DTT, 1 mM PMSF, 40 U/mL RNase Out (Invitrogen) pH 7.2) and slowly rocked for 5 min at 4°C. Plated was placed upright, buffer was drained to the bottom, and soluble cytosolic material was transferred to a chilled 1.5 mL microcentrifuge tube. Plate was gently washed with 5 mL of wash buffer (110 mM KOAc, 25 mM K-HEPES, 2.5 mM Mg(OAc)_2_, 1 mM EGTA, 0.004% digitonin, 1 mM DTT, 1 mM PMSF, pH 7.2) and buffer was subsequently discarded. Flask was coated with 0.5 mL lysis buffer (400 mM KOAc, 25 mM K-HEPES, 15 mM Mg (OAc)_2_, 1% NP-40, 0.5% DOC,1 mM DTT, 1 mM PMSF, 40 U/mL RNase Out, pH 7.2) and rocked for 5 min at 4°C. Plate was placed upright, buffer was drained to the bottom, and membrane-bound material was transferred to a chilled micro-centrifuge tube. Lysates were clarified at 7,500x g for 10 min at 4°C, and supernatants were transferred to new tubes. Fraction cleanliness was tested by Western blot, loading equal amounts of lysate (10 μg) for each fraction. Tubulin was used as a cytosolic (or soluble cytosolic) marker, RCC1 as a nuclear marker, and BiP as a membrane-associated marker.

### Western blots

Whole cell lysates were generated by lysing cells into buffer containing phosphatase inhibitors sodium fluoride (10 mM), β-glycerophosphate (1 mM) and sodium orthovanadate (1 mM, all Sigma). Protein samples were separated on 10,12, or 14% SDS-PAGE gels and transferred to nitrocellulose membranes (Bio-Rad, Hercules, CA) using a Trans-Blot Turbo Transfer System (Bio-Rad). Membranes were incubated for 1 hour at room temperature in blocking buffer (5% w/v non-fat dry milk in TBS + 0.1% Tween 20 (TBS-T)). Primary and secondary antibodies were incubated in either 5% w/v non-fat dry milk or 3% w/v bovine serum albumin (BSA), Research Products International, Mount Prospect, IL) in TBS-T, depending on manufacturer’s instructions. Dilutions used for primary antibodies: rabbit anti-β-tubulin (9F3) (1:1000; Cell Signaling Technology (CST), Danvers, MA), mouse anti-RCC1 (E-6) (1:1000; sc-55559, Santa Cruz Antibodies, Dallas, TX), rabbit anti-NRMT1 (1:1000; ^1^), rabbit anti-3me-SPK (1:10,000; ^6^), rabbit anti-1/2me-SPK (1:5000; ^6^), rabbit anti-2me-PPK (1:1000; ^1^), mouse anti-Importin-7 (E-2) (1:1000; sc-365231, Santa Cruz), mouse anti-ZHX2 (D-2) (1:1000; sc-393399, Santa Cruz), rabbit anti-BiP (C50B12) (1:1000, CST), rabbit anti-RPS25 (1:1000; Invitrogen), rabbit anti-RPL10A (1:1000; Bethyl Laboratories, Montgomery, TX), rabbit anti-eIF2α (D7D3) (1:1000, CST), rabbit anti-pSer51-eIF2α (D9G8) (1:1000, CST), rabbit anti-eIF3A (D51F4) (1:1000, CST). Secondary antibodies used were donkey anti-mouse or donkey anti-rabbit (1:5000; Jackson ImmunoResearch, West Grove, PA). Blots were developed on a ChemiDoc Touch imaging system (Bio-Rad) using Clarity Western ECL Substrate (Bio-Rad) or SuperSignal West Femto Maximum Sensitivity Substrate (Thermo Scientific).

### Mass Spectrometry

Twenty-four hours prior to transfection, 1×10^6^ HEK293T cells were plated in 10-cm tissue culture dishes. Cells were calcium phosphate transfected with 1 μg NRMT1-FLAG constructs and cultured for an additional 24 hours. Untransfected cells served as a control. Lysates from both untransfected and NRMT1-FLAG expressing cells were then fractionated to generate nuclear and cytoplasmic fractions and subsequently evaluated for purity as previously described. Multiple plates were used to generate sufficient material for the IP experiments. 3000 μg of pooled cytoplasmic lysate or 2000 μg of pooled nuclear lysate was then added to 20 μL of washed Pierce Protein G agarose beads (Thermo Fisher Scientific), and the mixture was rotated for 1 hr at 4°C to preclear lysates. After preclearing, the lysate was spun quickly, and the supernatant was added to 15 μL of washed anti-FLAG magnetic agarose beads (Thermo Fisher Scientific – Pierce). The mixture was rotated for 2 hr at 4°C, then washed 3x with buffer (PBS, 0.1% NP-40, 150 mM NaCl). Interacting proteins were eluted into FLAG buffer (25 mM HEPES pH 7.5, 100 mM NaCl, 1 μM PMSF, 0.1 mg.mL FLAG peptide (Anaspec, Fremont, CA)). Elutions were sent for label-free quantification mass spectrometry at the Cornell Institute of Biotechnology. Top interactors were those that had 100% abundance in NRMT1-FLAG samples compared to the untransfected control.

### Polysome profiles and sucrose gradient analysis

Prior to harvest, cells were pre-treated for 10 min in 37°C incubator with culture media supplemented with 100 μg/mL cycloheximide (CHX). All subsequent solutions also contained CHX. Cells were collected by trypsinization and spun at 200 x g for 3 min at 4°C. Cell pellet was washed with ice-cold PBS and pelleted again. Wash buffer was removed and cells were lysed into polysome lysis buffer (20 mM Tris pH 7.5, 150 mM NaCl, 15 mM MgCl_2_, 100 μg/mL CHX, 1mM DTT, 0.5% Triton-X-100, 0.1 mg/mL heparin, 8% glycerol, 20 U/mL Turbo DNase (Invitrogen), 200 U/mL RNAseOUT Inhibitor (Invitrogen), 0.5% DOC, 1 mM PMSF, 25 μg/mL Aprotinin, 25 μg/mL Leupeptin). Lysates were incubated on ice for 30 min with occasional vortexing, then clarified with sequential centrifugation for 5 min at 1,800 x g and 10,000 x g. The RNA concentration was determined for the clarified lysates, and ~ 150 to 600 μg was loaded onto 10-40% sucrose gradient (+CHX) and centrifuged at 39,000 rpm for 2 hr at 4°C in a SW41Ti swinging bucket rotor. Gradients were fractionated using a BioComp gradient fractionator with continuous A_260_ measurement. Fractions were TCA-precipitated, acetone washed and resuspended in 100 mM Tris pH 8.0, 8M Urea, and 1x sample buffer for subsequent Western blot analysis.

